# GLI transcriptional repression regulates tissue-specific enhancer activity in response to Hedgehog signaling

**DOI:** 10.1101/732024

**Authors:** Rachel K. Lex, Zhicheng Ji, Kristin N. Falkenstein, Weiqiang Zhou, Joanna L. Henry, Hongkai Ji, Steven A. Vokes

## Abstract

Transcriptional repression needs to be rapidly reversible during embryonic development. This extends to the Hedgehog pathway, which primarily serves to counter GLI repression by processing GLI proteins into transcriptional activators. In investigating the mechanisms underlying GLI repression, we find that a subset of these regions, termed HH-responsive enhancers, specifically loses acetylation in the absence of HH signaling. These regions are highly enriched around HH target genes and primarily drive HH-specific limb activity. They also retain H3K27ac enrichment in limb buds devoid of GLI activator and repressor, indicating that their activity is primarily regulated by GLI repression. The Polycomb repression complex is not active at most of these regions, suggesting it is not a major mechanism of GLI repression. We propose a model for tissue-specific enhancer activity in which an HDAC-associated GLI repression complex regulates target gene expression by altering the acetylation status at enhancers.

## INTRODUCTION

Transcriptional repressors employ distinct mechanisms for regulating gene expression. Long-term repression is accompanied by topological and biochemical changes to DNA that can lock down transcription within a cell lineage. In contrast, transient repression is rapidly reversible, providing a mechanism for controlling gene activation during the dynamic process of embryogenesis. This control is especially important for spatially restricting gene expression until signal transduction mechanisms alleviate repressor activity.

For example, the induction of HH signaling within the posterior half of the developing limb bud rapidly inhibits the formation of truncated GLI repressor proteins, instead promoting the production of full-length GLI transcriptional activators (Harfe et al. 2004). Consequently, GLI activators are found within the posterior limb bud in cells receiving the HH ligand while GLI repressors in the anterior limb bud spatially restrict the domains of HH target gene expression (Wang et al. 2000; Ahn and Joyner 2004). As a repressor-driven system, most transcriptional targets do not actually require GLI activator for transcription, but can be activated by loss of GLI repressor alone. This property of de-repression rather than activation is exemplified by *Shh^-/-^* limb buds (constitutive GLI repression, no GLI activation), which have a nearly complete absence of digits and a severe reduction in limb size. The phenotype is markedly improved in double mutants that lack most or all GLI activity (Litingtung et al. 2002; te Welscher et al. 2002; Wijgerde et al. 2002; Bai et al. 2004; Bowers et al. 2012). In particular, GLI de-repression is sufficient to activate most GLI target genes in the limb bud, suggesting that the primary role of the HH pathway is to alleviate GLI repression (Lewandowski et al. 2015). The labile nature of GLI repression represents a key mechanism for the dynamic transcriptional regulation of HH targets as HH induction rapidly inactivates GLI repression, resulting in transcription of targets within 4-9 hours of stimulation (Harfe et al. 2004; Panman et al. 2006).

The mechanism underlying GLI repression is unknown but could in principle function either by excluding GLI activator binding or by recruiting co-repressors (Wang et al. 2010). Although the former category provides an attractive model for how GLI proteins might interpret gradients of HH ligand (Falkenstein and Vokes 2014), it fails to account for the large number of GLI target genes that are fully activated upon de-repression in the absence of HH signaling.

Several GLI co-repressors have been identified in various contexts, including Atrophin (Zhang et al. 2013), Ski (Dai et al. 2002) and tissue-specific transcription factors (Oosterveen et al. 2012). Members of the BAF chromatin remodeling complex have also been shown to regulate GLI repressor activity; however, as these mutations also affect GLI-activation, it is unclear if they specifically mediate repressive activity or if they are essential for all GLI transcriptional response (Jagani et al. 2010; Zhan et al. 2011; Jeon and Seong 2016; Shi et al. 2016). Additionally, GLI activators have been shown to recruit the histone-demethylase JMJD3 to remove H3K27Me3, a mark that is placed by the Polycomb Repressor Complex 2 (PRC2), leading to subsequent activation (Shi et al. 2014; Weiner et al. 2016). Other studies have also suggested interactions between Polycomb repression and HH signaling in the limb bud (Wyngaarden et al. 2011; Deimling et al. 2018). Because mutations in these candidates are pleiotropic, it has been challenging to determine if they directly mediate GLI repression, a task compounded by the dual roles of GLI proteins as transcriptional activators and repressors.

We have used a genomic approach to determine if the chromatin modifications at GLI binding regions (GBRs) are altered in response to GLI repression. We hypothesized that GLI repressors regulate gene expression by inactivating enhancer activity.

Consistent with this, we find that GLI repression regulates enhancer activity through the de-acetylation of Histone H3K27. This repression occurs independently of Polycomb activity. Enhancers regulated in this fashion mark known GLI limb enhancers, are highly enriched around HH-responsive genes, and primarily drive tissue-specific enhancer activity within HH-specific domains. Collectively, the results suggest that GLI repressors inhibit gene expression by altering enhancer activity, providing an explanation for the labile nature of GLI repression.

## RESULTS

### A subset of GLI binding regions is epigenetically regulated by HH signaling

Since most HH targets can be activated by loss of GLI repression, we hypothesized that enhancers may be activated by HH signaling when GLI repression is relieved. To test this, we first identified active GLI enhancers in the developing limb at embryonic day 10.5 (E10.5), when high levels of HH target gene expression are observed. We used an endogenously FLAG tagged *Gli3* allele to identify GLI3 binding regions and then identified regions enriched for H3K27ac, a marker associated with active enhancers, by chromatin immunoprecipitation (ChIP-seq) (Heintzman et al. 2007; Heintzman et al. 2009; Creyghton et al. 2010; Rada-Iglesias et al. 2011; Cotney et al. 2012; Lopez-Rios et al. 2014; Lorberbaum et al. 2016). Altogether we identified 7,282 endogenous GLI3 binding regions (GBRs), with the majority of regions enriched for H3K27ac (83%) (Fig. 1A; Figure 1-Source Data 1).

**Figure 1.**
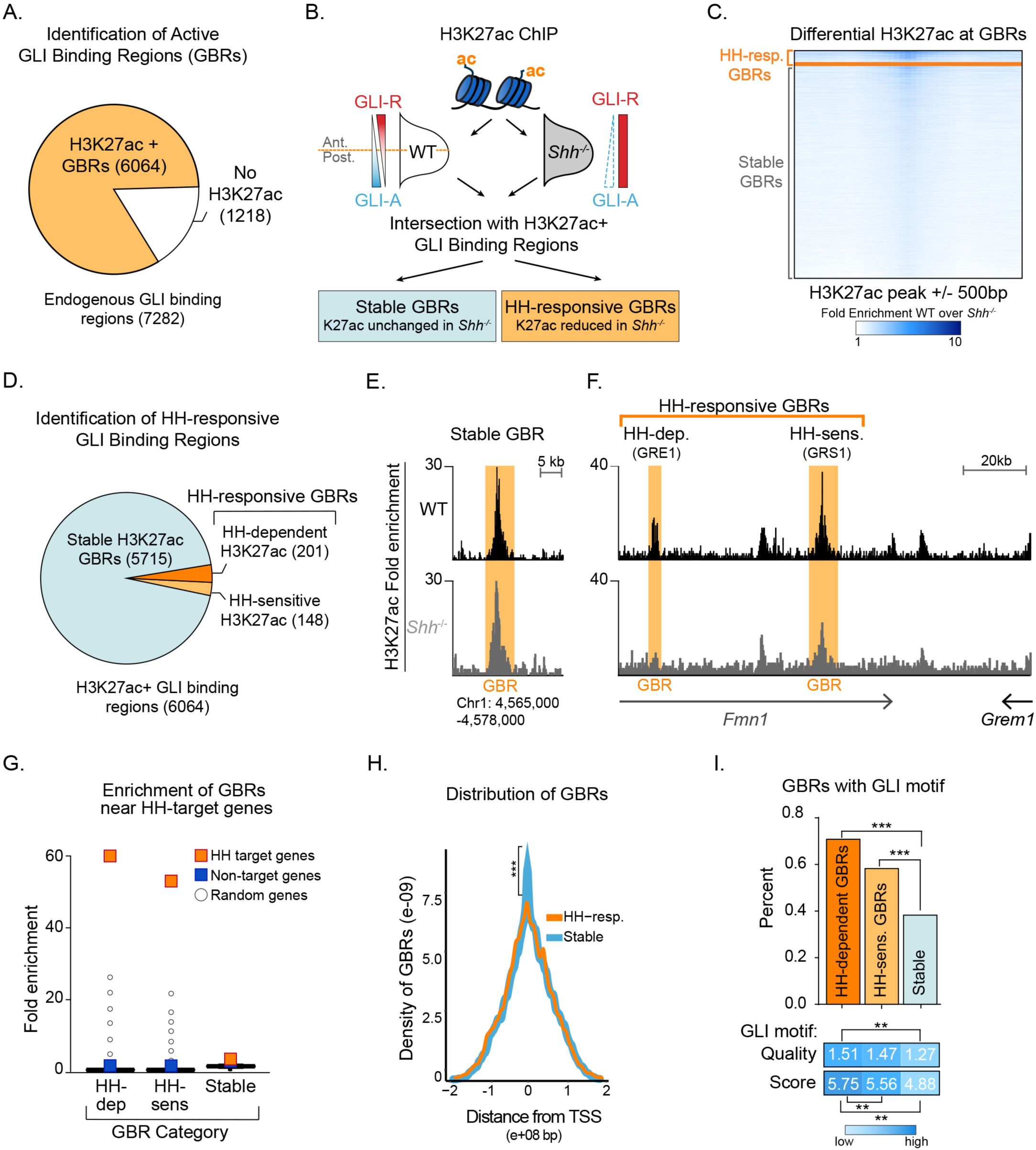
Hedgehog signaling regulates acetylation of H3K27 at a subset of GLI binding regions. A. Intersection of endogenous GLI3 and H3K27ac binding regions in WT E10.5 limb buds (n=2). B. Pipeline for identifying different categories of GLI bound regions. C. Heatmap depicting differential H3K27ac enrichment in WT over *Shh*^-/-^ limb buds for HH-responsive and Stable GBRs. D. Classification of GBR categories from E10.5 GBRs with H3K27ac in WT limbs. E-F. H3K27ac enrichment in WT and *Shh* ^-/-^ is shown across a representative genomic region near a Stable GBR (E) and two biologically validated HH-responsive GBRs: GRE1 (Li et al. 2014) and GRS1(Zuniga et al. 2012) at the HH target gene *Gremlin 1* (*Grem1*) (F). G. HH-dependent GBRs, HH-responsive GBRs and Stable GBRs are significantly enriched near HH target genes compared to randomly chosen genes (p=0 p=0, and p=0 respectively permutation test based on 10,000 permutations). H. Distribution of Stable and HH-responsive GBRs arounds transcription start sites (TSS), indicating significant enrichment of Stable GBRs (63%) at TSS compared to HH-responsive GBRs (26%) (p=2.55e-40, Fisher exact test, two sided). I. Both HH-dependent and HH-sensitive GBRs have significantly more GLI motifs than Stable GBRs (top)(p=2.2e-16 and p=8.00e-06; one-sided proportional test). GBRs containing GLI motifs have significantly more motifs per GBR within HH-dependent GBRs than Stable GBRs (p=5.92e-06; one-sided Wilcoxon test) and the quality of GLI motifs is significantly higher for HH-dependent and HH-sensitive GBRs than Stable GBRs (p= 5.03e-13 and p=5.98e-08; one-sided Wilcoxon test). See Figure1-Figure Supplement 1, Figure 1-Source Data 1, Figure 1-Source Data 2, Figure1-Source Data 3.

Next, we assessed changes in H3K27ac enriched GBRs in wild-type (WT) and *Sonic hedgehog* (*Shh*) null forelimbs. Since *Shh^-/-^* forelimbs have constitutive GLI repression, we hypothesized that in the absence of HH signaling, GLI repressor may prevent activation of its enhancers. We performed ChIP-seq for H3K27ac in E10.5 forelimbs, prior to overt phenotypes in *Shh* nulls (Chiang et al. 2001). Although most H3K27ac enriched regions were present in both WT and *Shh*^-/-^ embryos, a subset of 2,113 WT H3K27ac enriched regions had acetylation that was significantly reduced or completely lost in the absence of HH signaling (Figure 1-Source Data 2). We then asked whether those regions with reduced acetylation in the absence of HH signaling include GLI-bound enhancers. Most GBRs with H3K27ac enrichment in WT limbs retain H3K27ac in *Shh*^-/-^ limb buds, which we term Stable GBRs (Fig. 1C,D). This suggests that they function as active enhancers whose activity is not predominantly regulated by HH signaling. Interestingly, in a smaller subset of GBRs, H3K27ac enrichment was reduced or lost in the absence of HH signaling, suggesting that GLI repressor may regulate the activity of this group of enhancers. Within this GBR class with HH-responsive acetylation, we identified populations of GBRs that had either significant reductions (termed HH-sensitive) or a complete absence of H3K27ac enrichment (termed HH-dependent) in *Shh*^-/-^ limb buds (Fig. 1C,D). The latter two categories are henceforth collectively referred to as HH-responsive GBRs.

### Hedgehog-responsive GBRs are enriched near Hedgehog target genes

To determine if HH-responsive GBRs are associated with HH target genes, we examined biologically validated GLI enhancers in the *Ptch1* and *Gremlin* loci that mediate limb-specific transcription of these HH targets and found that they are among the HH-responsive class of GBRs (Fig. 1E) (Vokes et al. 2008; Zuniga et al. 2012; Li et al. 2014; Lopez-Rios et al. 2014). This suggests that HH-responsive enhancers may regulate limb-specific gene expression in response to HH signaling. Consistent with this possibility, we found that HH-responsive GBRs are highly enriched around genes that have reduced expression in *Shh*^-/-^ limb buds (Lewandowski et al. 2015). In contrast, Stable GBRs have minimal enrichment, however are still significantly enriched around both HH target and non-target genes (p=0, permutation test; Fig. 1G).

As many HH-responsive H3K27ac regions are not bound by GLI3, we asked if they clustered near GBRs. HH-responsive non-GLI binding regions are significantly clustered around HH-responsive GBRs, and to a lesser extent, near Stable GBRs (Figure 1-Figure Supplement 1B,C). We conclude that HH-responsive GBRs cluster with other HH-responsive regulatory regions, and are strongly associated with HH target genes, supporting their role in driving gene expression in response to HH signaling during limb development.

### HH-responsive GBRs are distal enhancers containing high quality GLI motifs

Although Stable GBRs are not highly enriched at HH target genes, 62% of them are located in close proximity to the promoters of genes, compared to 26% of HH-responsive GBRs (Fig. 1H). Most promoter-associated Stable GBRs (90%) are found at promoters with CpG islands, a quality typically associated with housekeeping genes and genes that tend to be more broadly expressed and less tissue-specific (Zhu et al. 2008). To examine how different classes of GBRs might be differentially regulated, we examined their GLI binding motifs. A higher percentage of HH-dependent and HH-sensitive GBRs contain GLI motifs compared to Stable GBRs. HH-dependent and HH-sensitive GBRs also contain a higher density and higher quality of GLI motifs compared to Stable GBRs (Fig. 1I). Interestingly, we did not uncover high levels of enrichment of other motifs using de novo motif analysis (Figure 1-Source Data 3). Taken together, we find that HH-responsive GBRs are distal elements containing a higher quality and density of GLI motifs compared to stable GBRs.

### The Polycomb Repressor Complex does not regulate most GLI enhancers

GLI activators recruit demethylases that remove H3K27me3, a hallmark of the Polycomb repressor complex (PRC2) to promote transcriptional activation of HH target genes, most notably *Gli1* and *Ptch1* (Margueron and Reinberg 2011; Shi et al. 2014; Lorberbaum et al. 2016). If PRC2 is recruited by GLI repressors, there should be enrichment of H3K27me3 at HH-responsive enhancers in *Shh^-/-^*, where maximal levels of GLI repression would lead to recruitment of PRC2 and thus methylation at these enhancers. Contrary to this prediction, we identified a minimal number of HH-responsive GBRs enriched for H3K27me3 in E10.5 *Shh*^-/-^ limb buds (31/349 GBRs; Fig. 2A-C, Figure 2-Source Data 1). As reported for MEFs (Shi et al. 2014), these methylated GBRs include the pathway target *Gli1* in addition to other pathway target genes such as *Ptch1* and *Ptch2* (Fig. 2B). In contrast, most HH-responsive GBRs and target gene promoters lack enrichment of H3K27me3 in the absence of HH signaling (Fig. 2C; Figure 2-Figure Supplement 1; Figure 2-Source Data 2). We conclude that while the PRC2 complex may play a role in the regulation of a small number of HH pathway target genes, it is not the primary mechanism by which GLI repressors inhibit developmental gene expression.

**Figure 2.**
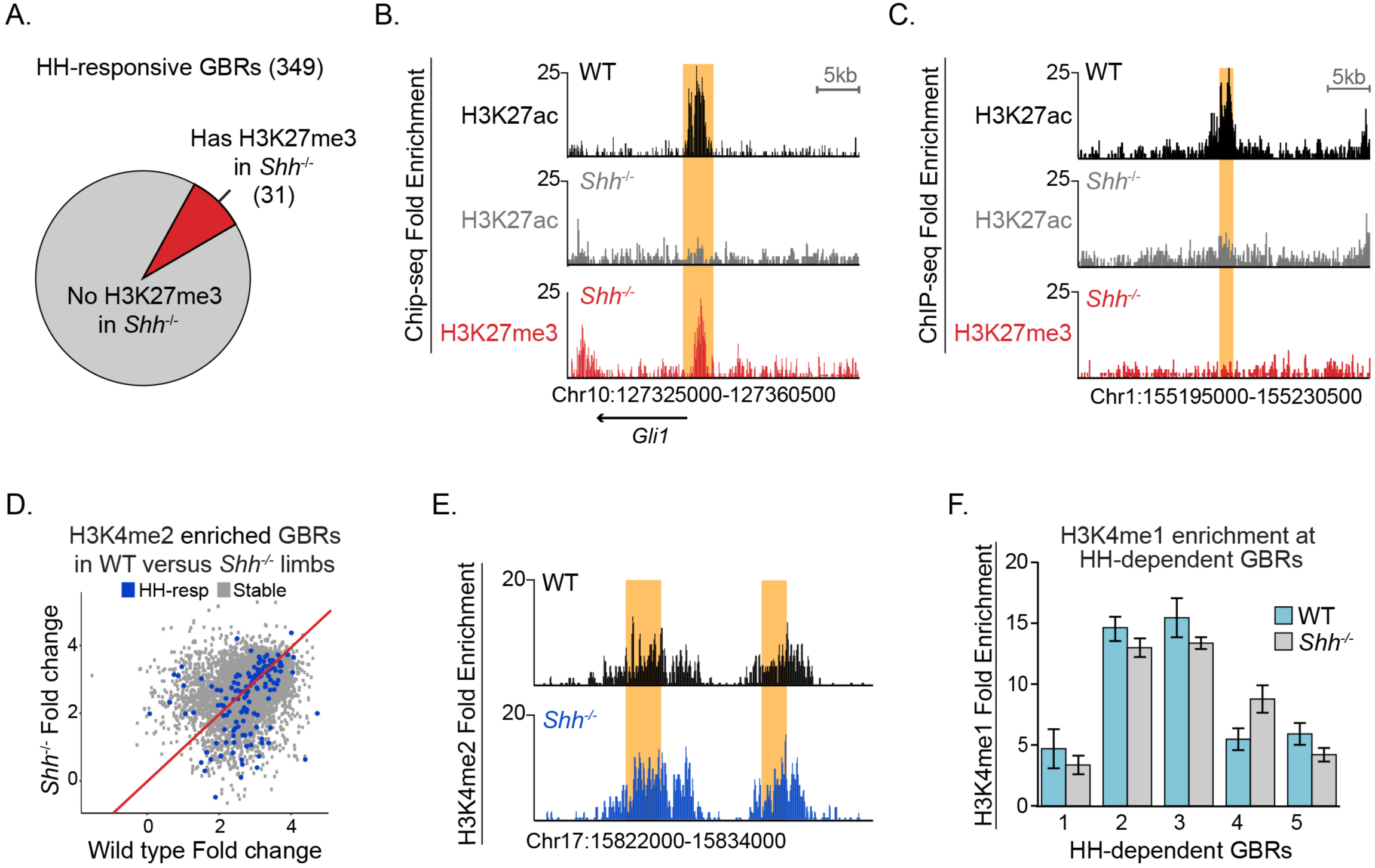
Most HH-responsive GBRs are not regulated by Polycomb repression and retain markers of poised enhancers. A. Chart depicts HH-responsive GBRs that contain enrichment for the PRC2 marker H3K27me3 in *Shh*^-/-^ limb buds (n=2). B. Tracks depicting a HH-responsive region in *Gli1* with differential H3K27ac enrichment in WT and *Shh*^-/-^ limb buds and H3K27me3 enrichment in *Shh*^-/-^ limb buds. C. Tracks depicting a representative HH-dependent GBR that also lacks H3K27me3. D. Scatter plot for H3K4me2 enrichment of Stable and HH-responsive GBRs from WT and *Shh*^-/-^ limb buds (n=2). No GBRs show significant changes in di-methylation of H3K4 between WT and *Shh^-/-^*. E. Representative track showing comparable levels of H3K4me2 enrichment for a HH-responsive GBR in WT and *Shh*^-/-^ limb buds. G. Quantitative-PCR assays indicating H3K4me1 ChIP enrichment in WT and *Shh*^-/-^ limb buds at HH-dependent GBRs (n=2). See Figure2-Figure Supplement 1, Figure2-Source Data1, Figure2-Source Data2, Figure 2-Source Data 3.

### Hedgehog signaling does not regulate other histone modifications at enhancers

We considered two possible mechanisms by which GLI repression could regulate H3K27ac enrichment in response to HH signaling: first, GLI repression could cause large-scale modifications to chromatin at enhancers resulting in an overall loss of their identity as enhancers. Alternatively, GLI repressors could regulate H3K27ac specifically. To address the first mechanism, we asked if HH regulates H3K4me2, another histone modification enriched at active or poised enhancers (Pekowska et al. 2011; Wang et al. 2014). H3K4me2 enriched at most GBRs in close proximity to promoters, being enriched at 26% of HH-responsive GBRs and 73% of Stable GBRs, none of which had significant reductions in H3K4me2 in *Shh*^-/-^ limbs compared to WT controls (Fig. 2D,E). Furthermore, essentially all peaks remained unchanged between the two genotypes, where only 12 peaks were reduced in *Shh^-/-^* limbs, none overlapping with GLI binding regions or non-GBR HH-responsive peaks (Figure 2-Source Data 3).

As H3K4me2 marked most Stable GBRs, but only a subset of HH-responsive GBRs, we also asked if H3K4me1, another modification associated with active and poised enhancers, was altered at HH-responsive GBRs in response to HH signaling (Heintzman et al. 2007; Heintzman et al. 2009; Creyghton et al. 2010; Rada-Iglesias et al. 2011). We performed ChIP on WT and *Shh*^-/-^ limb buds and assessed enrichment of H3K4me1 at several HH-responsive GBRs by quantitative PCR, selecting intergenic regions that would not overlap with promoters. All tested regions retained H3K4me1 enrichment in *Shh*^-/-^ limb buds (Fig. 2F). We conclude that HH-responsive regions retain enrichment of other active or poised enhancer marks, suggesting that HH signaling and GLI repression specifically regulate H3K27ac.

### Chromatin at HH-responsive GBRs compacts in the absence of Hedgehog

The dynamic acetylation of HH-responsive GBRs, yet unaltered methylation of H3K4 in in *Shh^-/-^* limb buds are properties consistent with ‘poised’ enhancers, which retain H3K4me1 and accessible chromatin in the absence of H3K27ac (Heintzman et al. 2009; Creyghton et al. 2010; Rada-Iglesias et al. 2011). Therefore, if HH-responsive enhancers are not active but ‘poised’ in the absence of HH, we predicted that chromatin accessibility would be unchanged in response to HH signaling. Using ATAC-seq to measure regions of open chromatin, we compared the accessibility of GBRs between WT and *Shh^-/-^* posterior limb buds, a fraction providing a more homogenous WT population of cells exposed to HH signaling (Fig. 3A; Figure 3-Source Data 1) (Buenrostro et al. 2013; Buenrostro et al. 2015). Overall, a higher percentage of Stable GBRs are accessible in WT limb buds than HH-responsive GBRs, suggesting a more restricted accessibility of HH-responsive GBRs even in WT conditions (Fig. 3B-C). Contrary to expectations for a poised enhancer, both HH-sensitive and HH-dependent GBRs have significantly reduced accessibility compared to Stable GBRs in the absence of HH signaling, with the majority of HH-responsive GBRs being more compact in *Shh^-/-^* compared to wild-type limbs (Fig. 3D-F). Overall, we conclude that HH-responsive GBRs are less accessible than Stable GBRs, with access being further restricted in *Shh^-/-^* limb buds, which have constitutive GLI repression.

**Figure 3.**
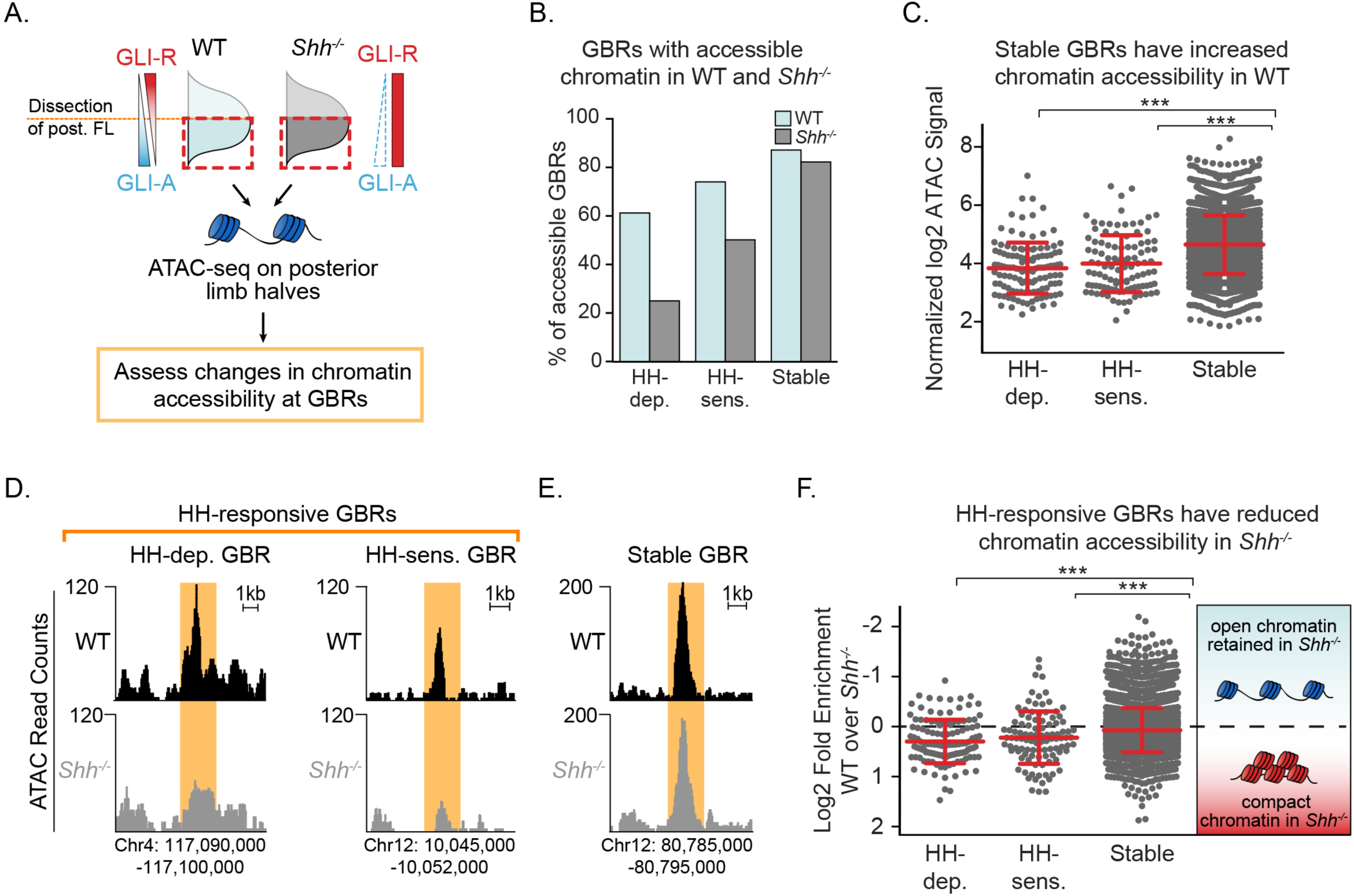
Chromatin accessibility is reduced in the absence of Hedgehog signaling. A. ATAC-seq pipeline for single pairs of dissected posterior halves of forelimbs (n=2). ATAC peaks, signifying accessible chromatin regions were intersected with Stable GBRs and HH-responsive GBRs. B. Many HH-responsive GBRs that are accessible in WT limb buds are inaccessible *Shh*^-/-^ limb buds, while the accessibility of Stable GBRs remain largely unchanged. C. Plot of log2 fold changes in chromatin accessibility in WT limbs indicating that Stable GBRs are more accessible than HH-dependent and HH-responsive GBRs (p= 3.98e-19, p= 9.21e-11; Wilcoxon rank sum test). Each data point represents a single GBR and red bars indicate the median, upper and lower quartiles. D-E. Representative ATAC-seq peaks showing lack of accessibility in *Shh*^-/-^ limb buds at HH-responsive GBRs (D), but not in Stable GBRs (E). F. Plot of log2 fold changes in chromatin accessibility in the presence and absence of HH signaling. HH-responsive GBRs are significantly less accessible than Stable GBRs (Stable vs. HH-sensitive. p=0.001; Stable vs. HH-dependent p= 4.99e-09; Wilcoxon rank sum test). See Figure3-Source Data 1.

### De-repression is the dominant mechanism regulating GLI enhancer activation

The presence of multiple GLI proteins and their bifunctional roles as both transcriptional activators and repressors has made it challenging to determine how HH genes are primarily regulated. To test this on enhancers, we performed H3K27ac ChIP in *Shh^-/-^;Gli3^-/-^* limb buds (devoid of GLI activators and most GLI repressors). We hypothesized that loss of H3K27ac at most HH-responsive enhancers is due to constitutive GLI repression preventing acetylation of GLI enhancers. Thus, in *Shh^-/-^;Gli3^-/-^* limbs, we predicted H3K27ac should be maintained at HH-responsive enhancers. Alternatively, if GLI activator is required, H3K27ac would remain absent or reduced as it does in *Shh^-/-^* limbs.

Since *Shh^-/-^;Gli3^-/-^* mutants are phenotypically identical to *Gli3^-/-^* mutants and require genotyping and our ChIP protocol works best with fresh tissue, individual pairs of E10.5 limb buds (∼100k cells/pair) were processed independently to confirm genotypes. To overcome the reduced tissue available for ChIP samples, we optimized a ‘MicroChIP’ approach to allow ChIP-seq on single pairs of limb buds and assessed H3K27ac enrichment at GLI enhancers in *Shh^-/-^;Gli3^-/-^* limb buds (Fig. 4A; Figure 4-Source Data 1). As anticipated, we did have reduced signal, however we were still able to detect 60% of our HH-responsive GBRs and 91% of Stable GBRs. Consistent with expectations, HH-responsive GBRs associated with *Gli1* and *Ptch1*, which require GLI activation (Litingtung et al. 2002; te Welscher et al. 2002), had greatly reduced expression in the double mutants (Fig. 4B,D). A small number of additional GBRs were also reduced in double mutants, suggesting they too require GLI activation (Fig. 4D,E). However, consistent with a GLI repression-driven model, most HH-responsive GBRs retained or increased H3K27ac enrichment in the absence of both GLI activator and repressor (Fig. 4C-E). Despite being unchanged in *Shh^-/-^* limbs, Stable GBRs had slight but significant increases in H3K27ac enrichment (Fig. 4F), indicating that on a population level, some of these regions respond to GLI repression (see Discussion).

**Figure 4.**
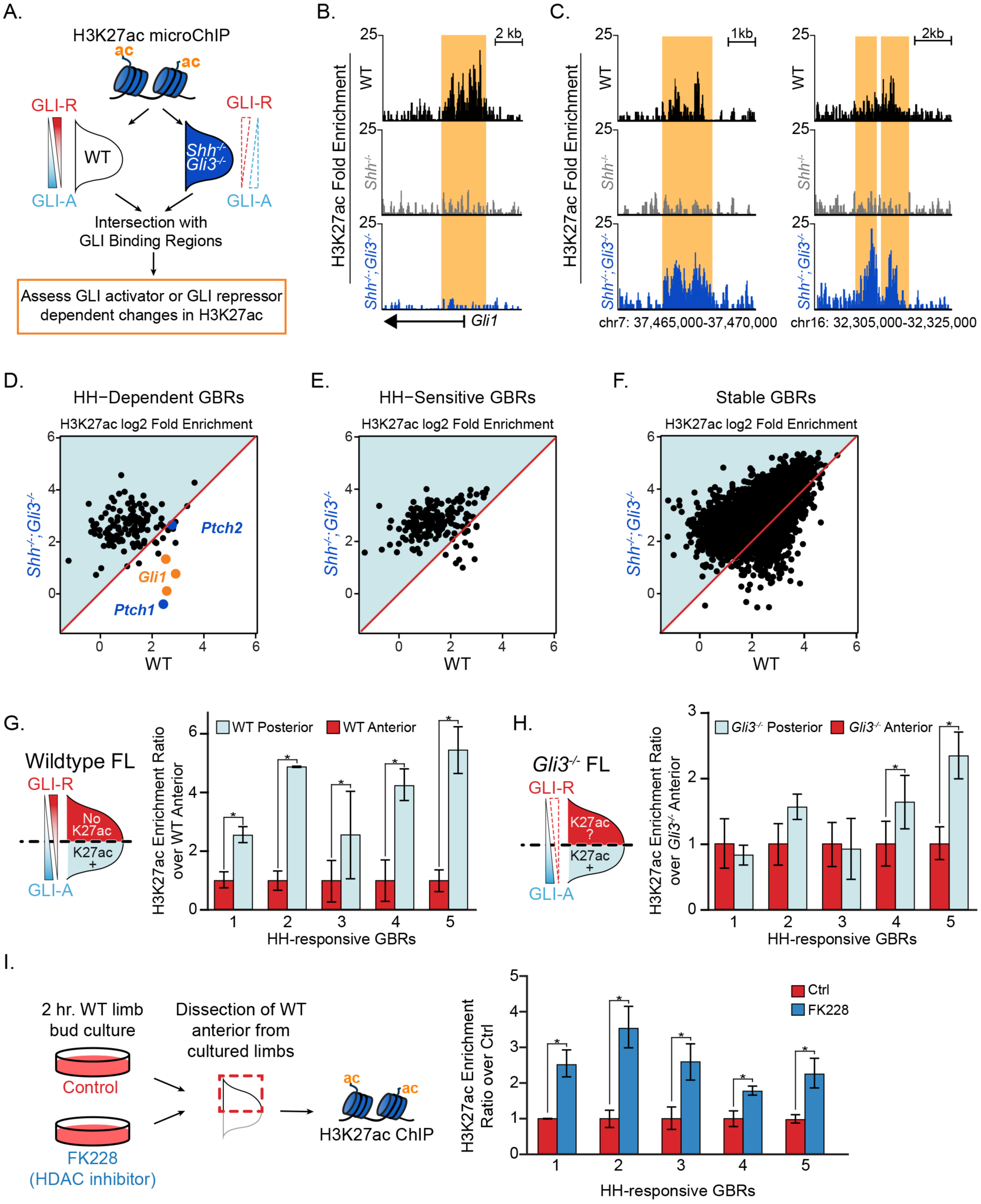
GLI de-repression activates most HH-responsive enhancers. A. *Shh^-/^;Gli3^-/-^* H3K27ac ‘MicroChIPs’ on single pairs of E10.5 forelimbs (33-34S) *Shh^-/-^;Gli3^-/-^* and WT littermate controls (n=2, respectively). B. A HH-responsive GBR near *Gli1* which requires GLI activator for H3K27ac enrichment. C. Representative examples of HH-responsive GBRs, activated by loss of GLI repressor that do not require GLI activator. D-F. Scatter plot of H3K27ac enrichment of HH-dependent, HH-sensitive and Stable GBRs in WT and *Shh^-/-^;Gli3^-/-^* limbs. Each dot represents a single GBR. The p-values indicate a significant enrichment of acetylation in *Shh^-/-^;Gli3^-/-^* among all GBR classes (Wilcoxon-rank sum tests). G-H. E10.5 WT and *Gli3^-/-^* limb buds were dissected into anterior and posterior halves as indicated and selected HH-dependent GBRs were tested for H3K27ac enrichment by quantitative PCR in each fraction (n=4). HH-dependent GBRs have higher ratios of posterior to anterior H3K27ac enrichment in WT limb buds (G) while most HH-dependent GBRs have equal ratios of posterior to anterior H3K27ac enrichment in *Gli3^-/-^* limb buds (H) (asterisks indicate p<0.05, paired T-test). I. Inhibition of HDAC 1/2 using 250nM of FK228 in cultured limb buds for two hours resulted in significant increases of H3K27ac enrichment in anterior cultured limb buds compared to control anterior limbs (n=4; asterisks indicate p<0.05, paired T-test). See Figure 4-Figure Supplement 1, Figure 4-Source Data 1.

In a parallel series of experiments, we noted that HH-responsive GBRs have higher levels of H3K27ac enrichment in posterior limb halves compared to anterior limb halves (Fig. 4G). This contrasts with *Gli3^-/-^* limb buds, where H3K27ac levels in anterior halves are comparable to those in posterior halves (Fig. 4H), a finding that is also consistent with a GLI repressor driven model. Together these results strongly support a GLI repressor centric mode of regulation where GLI de-repression is responsible for activation of most GLI limb enhancers. We conclude that GLI activator does not mediate acetylation levels at most HH-responsive GBRs.

### HDAC1 is responsible for loss of H3K27ac at HH-responsive enhancers

The above results support a model in which loss of an HDAC-GLI repressor complex leads to acetylation. To test this, we cultured limb buds in the presence of the HDAC1/2 inhibitor FK228 (Furumai et al. 2002), yielding greatly upregulated levels of H3K27ac within two hours of treatment (Figure 4-Figure Supplement 1). We then dissected the anterior halves of limb buds cultured in control or FK228-containing media and compared the levels of H3K27ac enrichment at HH-responsive GBRs shown to have enriched H3K27ac levels in posterior limb halves. Inhibition of HDACs resulted in increased acetylation at HH-responsive enhancers compared to untreated control anterior limb buds (Fig. 4I). We conclude that GLI repressors regulate H3K27ac levels at HH-responsive GBRs at least in part through HDAC1/2.

### HH-responsive GBRs have increased tissue-specificity compared to Stable GBRs

Having identified distinct classes of GBRs that respond differently to HH signaling, we next addressed the biological significance of this enhancer behavior. To this end, we used the VISTA enhancer database to identify a total of 305 Stable and 23 HH-responsive GBRs that had been tested for enhancer activity (Visel et al. 2007). While nearly half of each class have enhancer activity in the limb, HH-responsive GBRs tend to have activity specific to the HH-responsive posterior limb bud while Stable GBRs tend to have activity throughout the limb or regions that are not responsive to HH (Fig. 5A,B) (Ahn and Joyner 2004; Probst et al. 2011; Lewandowski et al. 2015). Furthermore, HH-responsive enhancers are active more specifically within the limb (an average of 1.9 tissues) while Stable GBRs drive expression in more tissues (an average of 2.9 tissues; P<0.01; Fig. 5C; Figure 5-Source Data 1). These results point to a role for Stable GBRs as general enhancers that drive expression in multiple tissues, while HH-responsive GBRs mediate tissue-specific expression.

**Figure 5.**
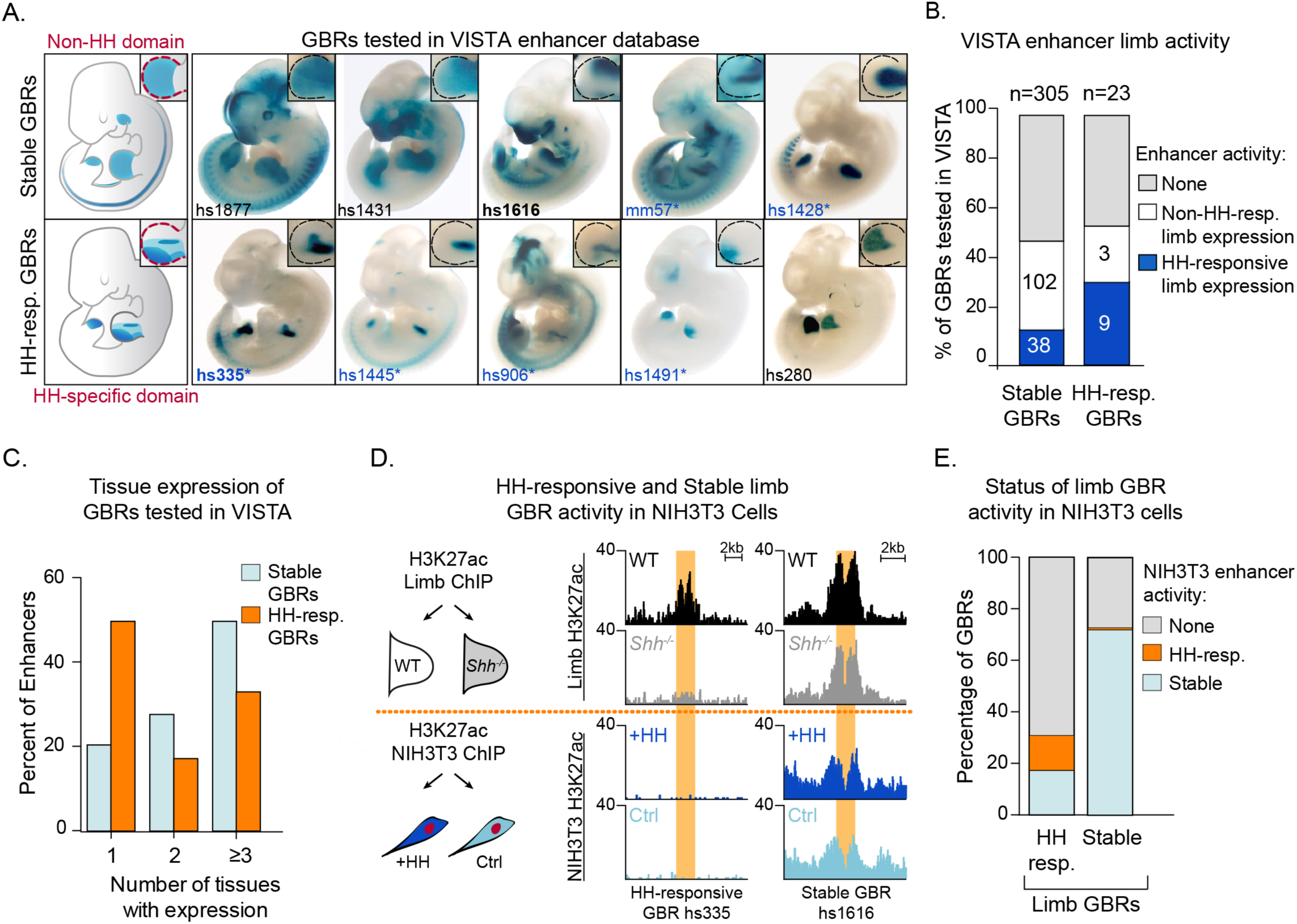
Hedgehog-responsive GBRs have tissue-specific enhancer activity within HH-specific domains. A. Enhancers with annotated limb activity in VISTA corresponding to representative HH-responsive GBRs (bottom) and Stable GBRs (top) with limbs magnified and outlined in insets. Limb buds containing HH-specific domains of enhancer activity are indicated by an asterisk. B. Chart indicating total number of VISTA enhancers tested for HH-responsive and Stable GBRs, the numbers of enhancers for each category and their limb enhancer activity. C. Chart delineating the percentage of HH-responsive and Stable limb enhancers that drive expression in one or more tissues. D. Schematic of NIH3T3 H3K27ac ChIP treated with and without the HH agonist purmorphamine (+HH) and the activity of representative HH-responsive and Stable limb GBRs in response to HH activation in limb and NIH3T3 cells (n=2). E. Graph indicating how the acetylation status of HH-responsive and Stable limb GBRs responds to HH signaling in HH-responsive NIH3T3 cells. See Figure 5-Source Data1, Figure 5-Source Data 2.

To examine this more systematically, we treated HH-responsive NIH3T3 cells with and without the HH agonist purmorphamine, identified H3K27ac enriched regions by ChIP-Seq, and assessed the H3K27 acetylation status of different classes of limb GBRs. Strikingly, only 12% of HH-responsive limb GBRs are also HH-responsive in NIH3T3 cells. An additional 18% of HH-responsive limb enhancers have stable acetylation in NIH3T3 cells, while most lack any activity. In contrast, 70% of Stable GBRs in the limb are still active in NIH3T3 cells (Fig. 5D,E; Figure 5-Source Data 2). We conclude HH-responsive GBRs are tissue specific enhancers that mediate HH signaling while Stable GBRs have broadly expressed enhancer activity.

## DISCUSSION

We find that a subset of GLI-bound enhancers has chromatin modifications that change in response to HH signaling. Compared to WT embryos, these regions have reduced or absent levels of histone H3K27 acetylation in *Shh^-/-^* embryos, suggesting a loss of enhancer activity. Many previously validated GLI limb enhancers are HH-responsive, including those regulating *Grem1*, *Ptch1* and *Gli1* (Fig. 1G) (Vokes et al. 2008; Zuniga et al. 2012; Li et al. 2014; Lopez-Rios et al. 2014). Moreover, HH-responsive GBRs are highly enriched near HH target genes while the much larger class of Stable GBRs are not (Fig. 1E). This suggests that HH target gene regulation is primarily mediated through HH-responsive GBRs. The discovery of this response provides important information about the mechanism of GLI repression. It also provides a predictive tool for identifying enhancers regulating HH target genes in other biological contexts.

We propose a model in which GLI repression primarily regulates enhancer activity by deacetylation of histone H3K27. Because H3K4me1 and H3K4me2 levels are unchanged during maximal GLI repression, these enhancers presumably remain poised for activation, albeit in a less accessible state. Upon binding HH-responsive enhancers, GLI repressors recruit HDACs, which prevent otherwise competent enhancers from acquiring enriched H3K27 acetylation. The loss of GLI repression, either genetically (*Shh^-/-^;Gli3^-/-^* or *Gli3^-/-^* limb buds), or developmentally (initiation of *Shh* expression) results in a loss of GLI repression and accompanying HDAC activity (Fig. 6A). This chromatin-based mode of regulation enables the dynamic control of a field of cells containing primed enhancers. To determine if this priming event occurs on an *ad hoc* basis by disparate inputs or if it is the result of coordinated, HH-independent signaling events, we examined HH-responsive GBRs for the enrichment of additional binding motifs. Besides the GLI motif itself, no other motifs are enriched at high levels (Figure 1-Source Data 3) suggesting that HH-responsive GBRs are a heterogenous population of enhancers with no predominant co-regulators.

**Figure 6.**
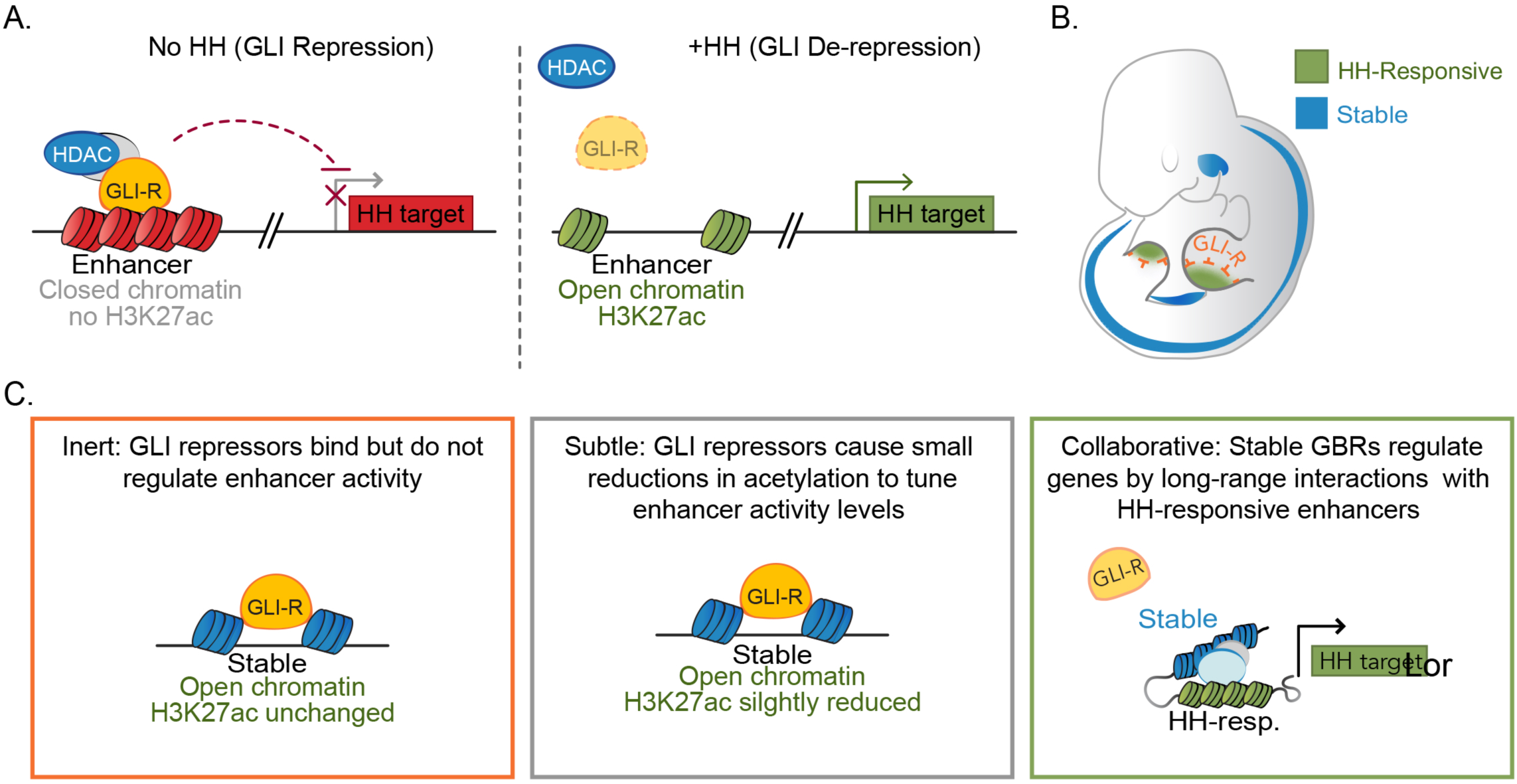
Model for GLI transcriptional repression. A. In the absence of HH, GLI repressors bind to enhancers for HH target genes, limiting their accessibility and recruiting an HDAC complex that de-acetylates Histone H3K27, inactivating the enhancer. In the presence of HH signaling, GLI de-repression and loss of associated HDAC activity result in increased accessibility, the accumulation of H3K27ac and gene transcription. B. Schematic showing tissue-restricted activity of HH-responsive GBRs within HH-responsive gene expression domains. C. Possible roles for Stable GBRs in HH transcriptional regulation.

Despite being critical for the transcriptional regulation of HH targets, HH-responsive enhancers are a distinct minority, constituting 6% of all GLI-bound, active enhancers. The rest are Stable GBRs with an unclear role in HH transcriptional regulation. Although these enhancers do not have significantly reduced levels of H3K27 enrichment in *Shh^-/-^* limbs, some of them show a trend toward reduced H3K27ac that suggests a continuum of GLI-bound enhancers that range from completely HH-responsive (HH-dependent) to those Stable GBRs that have no HH response. Consistent with this, Stable GBRs do have a modest overall increase in H3K27ac enrichment in *Shh^-/-^;Gli3^-/-^* limbs on a population level, indicating that their H3K27ac levels are regulated by GLI repressor to some extent. On the other hand, these enhancers are enriched at CpG rich promoters, which are associated with more broadly expressed genes and are not enriched near HH target genes (Fig. 1G,H). They are also more highly conserved than HH-responsive GBRs (Figure 1-Figure Supplement 1D). In contrast to HH-responsive enhancers, they appear to be active in other cell types and tissues besides the limb (Fig. 6B). One possibility is that many Stable GBRs do not have a major role in mediating Hedgehog signaling; GLI repressors at these regions are relatively inert. A second possibility is that GLI repression at Stable GBRs mediates subtle changes to acetylation that confer small reductions in transcription that are beyond the limits of our detection. Finally, it is possible Stable enhancers are globally active, but engage in long-range collaborations with tissue specific HH-responsive enhancers to activate transcription (Fig. 6C).

Previous modeling has suggested that GLI repressors within an enhancer work cooperatively through multiple GLI sites (Parker et al. 2011), providing another mechanism for tuning enhancer response. HH responsive GBRs contain more GLI motifs than Stable GBRs, which may make them more responsive to GLI repression, although in contrast to those models, they have high quality GLI motifs. As many GLI target genes, including *Ptch1* and *Grem1,* are regulated by multiple GLI enhancers (Vokes et al. 2008; Zuniga et al. 2012; Li et al. 2014; Lopez-Rios et al. 2014; Lorberbaum et al. 2016), this integration likely extends to higher level hubs of enhancer organization. For example, HH-responsive H3K27ac regions that are not bound by GLI cluster near HH-responsive GBRs, as do Stable GBRs suggesting that they may be modified based on proximity to GLI-repressor-HDAC complexes (Figure 1-Figure Supplement 1B).

The majority of HH-responsive GBRs do not have H3K27me3 enrichment even when there is maximal GLI repression (Fig. 2A-D). This indicates that the Polycomb repressor complex is not involved in mediating most GLI repression, a conclusion that seemingly conflicts with several studies showing direct or indirect roles for PRC2 in repressing HH transcription. However, these studies largely considered the transcriptional activator targets *Ptch1* or *Gli1* or looked at genetic interactions (Wyngaarden et al. 2011; Shi et al. 2014; Lorberbaum et al. 2016; Shi et al. 2016; Deimling et al. 2018). Consistent with their findings, *Gli1* has high levels of H3K27me3 enrichment in *Shh^-/-^* limb buds (Fig. 2B). Although *Gli1* and *Ptch1* are often examined in the context of GLI de-repression, they are both GLI-activator genes in that they require the loss of GLI repression as well as subsequent GLI activation for their expression (Litingtung et al. 2002; te Welscher et al. 2002). GLI activator targets such as these are likely to differ fundamentally in their mode of regulation from those that are activated upon de-repression. As H3K27me3 enrichment is commonly found at promoters (Young et al. 2011), GLI repressors on distal enhancers not directly enriched by H3K27me3 might still facilitate the recruitment of PRC2 to promoters through enhancer-promoter interactions. However, only one third of Hedgehog target genes have H3K27me3 enrichment at their promoters (Figure 2-Figure Supplement 1; Figure 2-Source Data 2), arguing against this scenario. Interestingly, HH-responsive GBRs enriched for H3K27me3 in *Shh^-/-^* limbs have increased H3K27ac enrichment in *Shh^-/-^;Gli3^-/-^* limbs at all of these GBRs except for ones near the HH pathway genes *Gli1, Ptch1* and *Ptch2* (Fig. 4B). Thus, for rare GBRs requiring GLI activation, their mode of action is consistent with previously proposed models in which GLI activators recruit a complex to remove H3K27Me3, resulting in the activation of these enhancers and subsequently their cognate target genes (Shi et al. 2014).

Confusingly, HDACs have been shown to have properties both consistent with and contradictory to our model. HDACs bind to and deacetylate GLI1 and GLI2 proteins, promoting their ability to act as transcriptional activators (Canettieri et al. 2010; Coni et al. 2013; Mirza et al. 2019). HDACs have also been shown to bind cis-regulatory regions in *Gli1*, consistent with an additional role in positively regulating HH-mediated transcription (Zhan et al. 2011). On the other hand, a SKI-HDAC complex has been shown to bind to and interact genetically with GLI3 to repress anterior digit formation in the limb bud (Dai et al. 2002). Similarly, Atrophin acts as a GLI co-repressor by recruiting an HDAC complex (Zhang et al. 2013). Multiple studies with SWI/SNF BAF complex members also indicate that they regulate aspects of both GLI activation and repression, roles that have in some cases been shown to be directed by the dynamic association of BAF members with HDAC complexes (Jagani et al. 2010; Zhan et al. 2011; Jeon and Seong 2016). Collectively, these studies highlight the complexity of GLI regulation and the need for further studies to determine which complexes directly impact GLI repression.

## MATERIALS AND METHODS

### Embryonic manipulations

Experiments involving mice were approved by the Institutional Animal Care and Use Committee at the University of Texas at Austin (protocol AUP-2016-00255). The *Gli3^Xt-J^* and *Shh^tm1amc^* null alleles have been described previously (Hui and Joyner 1993; Dassule et al. 2000) and were maintained on a Swiss Webster background. The *Gli3^3XFLAG^* allele, with an N-terminal 3XFLAG-epitope, (Lopez-Rios et al. 2014; Lorberbaum et al. 2016) was maintained on a mixed background. For ChIP and ChIP-seq experiments, fresh E10.5 (32-35 somite) forelimb buds were pooled from multiple litters to obtain sufficient *Gli3*^-/-^ and *Shh*^-/-^ mutant embryos along with somite matched controls (Swiss Webster embryos for *Gli3*^-/-^ experiments and a mixture of WT and heterozygous littermates for *Shh*^-/-^) embryos. For ATAC-seq, fresh pairs E10.5 (35 somite) posterior forelimb buds were dissected from individual embryos.

To inhibit HDAC1/2, E10.5 embryos (32-35S) were dissected in warm limb bud culture media (Panman et al. 2006) and explants still attached to the body wall were cultured in 250nM of HDAC inhibitor FK228 (Selleckchem S3020), or DMSO vehicle control, for two hours at 37C. For each condition, 20-25 embryos were used (n=4). After incubation, the explants were changed into fresh media (without inhibitor) to dissect anterior limb buds. Cells from anterior limbs were then dissociated and processed for ChIP.

### Cell Culture

NIH3T3 cells were seeded on 6 cm plates with 5×10^5 cells and grown for three days until completely confluent. They were then switched to low serum (0.5%) and treated with 400nM purmorphamine (Stemgent 04-0009) or 0.01% DMSO (vehicle control) for 2 days.

### Western Blots

Whole limb buds from a single litter were lysed for 1 hour at 4C. For fractionation, 500,000 cells from limb buds were then dissociated with 100ug/mL Liberase (Roche 05401119001), resuspended in CSKT buffer (10mM PIPES pH6.8, 100mM NaCl, 300mM sucrose, 3mM MgCl_2_, 1mM EDTA, 1mM DTT, 0.5% TritonX-100, incubated on ice for 10 min, and centrifuged for 5 min @ 5000g. The cytoplasmic fraction (supernatant) and nuclear pellet were each resuspended in loading dye and boiled. Western blots were incubated with the following primary antibodies for 1 hour at room temperature in 3% milk: 1:4000 M2 Flag (Sigma 3165),1:4000 H3 (Cell Signaling 4499), 1:1000 GAPDH (Cell Signaling 5174), 1:1000 H3K27ac (Abcam Ab4729), 1:2000 B-actin (Cell Signaling 8457). Secondary antibodies were incubated for 1 hours at room temperature in 3% milk: 1:5000 Donkey anti-mouse (Jackson 715-035-150), Donkey anti-rabbit (Jackson 711-005-0152).

### Chromatin Immunoprecipitation

ChIP experiments were performed as previously described (Vokes et al. 2008) with the following modifications. Histones ChIPs were performed on whole E10.5 (32S-35S) forelimbs pooled from 6-8 embryos. The GLI3-FLAG ChIP and the H3K27ac ChIP on cultured and treated limbs were performed on E10.5 (32-35S) forelimbs from 20-25 pooled embryos. Cells were dissociated with 100ug/ml Liberase (Roche 05401119001) and fixed for 15 minutes at room temperature in 1% formaldehyde. After cell lysis, chromatin was sheared in buffer containing 0.25% SDS with a Covaris S2 focused ultrasonicator using the following settings: Duty Cycle: 2%, Intensity: 3, Cycles/burst: 200, Cycle time: 60 sec, Power mode: frequency sweeping. Sheared chromatin was then split into 3 ChIP reactions and incubated with antibody-dynabead preparations overnight. The H3K27ac antibodies for conventional ChIP were from Diagenode (C15200184) and Abcam (ab4729), while the H3K27Ac antibody for MicroChIPs was from Diagenode (C15410196). Additional antibodies recognized H3K4me1 (Millipore ABE1353) H3K4me2 (Millipore 07-030) and H3K27me3 (Abcam (ab7028). Beads were washed 5 times with RIPA buffer (1% NP40, 0.7% Sodium Deoxycholate, 1mM EDTA pH8, 50mM Hepes-KOH pH7.5, 2% w/v Lithium Chloride) and 1 time with 100mM Tris, 8.0, 10mM EDTA, 8.0, 50mM NaCl and then eluted at 70°C for 15 minutes. Crosslinking was reversed overnight at 70°C. Chromatin was purified and concentrated, then subjected to quantitative PCR and/or library preparation and sequencing. Quantitative PCR-based analysis was performed using SensiFAST SYBR-LoROX (Bioline BIO-94020) on a Viia7 system (Applied Biosystems). ChIP regions subsequently tested by qPCR are referred to in the figures by the unique peak ID number (Figure 1-Source Data 2). For each biological replicate, 2-3 technical replicates were performed for each qPCR reaction and the Ct values were averaged. Chromatin enrichment was determined by calculating delta delta Ct method (Livak and Schmittgen 2001) against a control region (C1).

Primers are described below. Primers are identified by their H3K27ac Peak ID. Primers labeled #1-5 are HH-dependent GBRs.

**45402 F** (B-actin normalizing primer): AGAAGGACTCCTATGTGGGTG

**45402 R** (B-actin normalizing primer): ACTGACCTGGGTCATCTTTTCA

**45402 F** (B-actin normalizing primer for H3K4me2): AGCTAACAGCCTGCCCTCTG

**45402 R** (B-actin normalizing primer for H3K4me2): TTTTCCGGTGGTACCCTAC

**NONE F** (Negative normalizing primer): GCCAGAATTCCATCCCACTA

**NONE R** (Negative normalizing primer): CCAATAACCTGCCCTGACAT

**32467 F** (#1): ACGCAGGCAGTTCCAATACA

**32467 R** (#1): AGGGACTTCACCCAGTTCCA

**15198 F** (#2): CCCTCCATTCTCCCTCCTTA

**15198 R** (#2): GGACCTTTCCGTTGAAGTGA

**2666 F** (#3): CTGGCTCCCAGAATCTCTCA

**2666 F** (#3): TTGTGCCCCATCTCTTTCAG

**45094 F** (#4): GGGAGGGGTGAACTTGTCTT

**45094 R** (#4): TGCAAATGAACACACGCATA

**20941 F** (#5): TTCCCAGCTCAAGGTCATGT

**20941 R** (#5): AGGAGGCAATGAAGACACTGG

Samples were processed for ‘MicroChIP’ using the Diagenode True MicroChIP kit (Cat #C01010130) with the following modifications. Briefly, individual limb pairs (∼100k cells) of wildtype, *Shh^-/-^* and *Shh^-/-^;Gli3^-/-^* E10.5 embryos (33-34S) were processed separately by dissociating limb buds with 100ug/mL Liberase (Roche 05401119001), crosslinked for 10 minutes, lysed and then sheared. Samples were sheared on a Diagenode BioRuptor for 6 cycles on high, 30sec on/off and processed through shearing while genotyping in parallel for *Shh^-/-^;Gli3^-/-^* and wildtype littermates (*Shh^+/+^;Gli3^+/+^*). Sheared chromatin was then incubated with H3K27ac antibody (Diagenode C15410196) overnight and Protein A magnetic beads (Diagenode C03010020) the following day for 2 hours. Chromatin-bound beads were washed, eluted and de-crosslinked and purified using MicroChIP DiaPure columns (Diagenode C03040001).

### ChIP-Seq

The ChIP-seq raw datasets from this study have been deposited in GEO (GSE108880) (see Source Data for Figures 1-5 for processed ChIP-seq and ATAC-seq data). All chromosomal coordinates refer to the mm10 version of the mouse genome. After ChIP was performed as described above, libraries were generated using the NEBNext Ultra II library preparation kit with 15 cycles of PCR amplification (NEB E7645). The libraries for the ‘MicroChIP’ samples were generated using the MicroPlex library prep kit (Diagenode C05010012) and sequenced to a depth of >40 million reads per sample for both ChIP and ‘MicroChIP’ experiments, using two biological replicates. Peaks were called using CisGenome version 2.1.0 (Ji et al. 2008). To identify differentially enriched peaks in the WT and *Shh^-/-^* limb buds (or control and purmorphamine-treated NIH3T3 cells), the peaks were merged to determine how many WT, WT input, *Shh^-/-^* and *Shh^-/-^* input reads overlapped with the peak region. The read numbers were adjusted by library size and log2 transformed after adding a pseudo-count of 1. The differential analysis between WT and WT input used limma (Ritchie et al. 2015). The FDR of the differential test was obtained and peaks with FDR < 0.05 are determined as having differential signal between WT and WT input. The same differential analysis procedure was repeated to compare between *Shh^-/-^* and *Shh^-/-^* input, and between WT and *Shh^-/-^*. To determine GLI motif quality, de novo motif discovery was performed on the 1000 GBRs with the highest quality using the flexmodule_motif function in CisGenome to identify the GLI motif. The GLI motif was mapped to the mouse genome using the motifmap_matrixscan_genome function in CisGenome software with default parameters.

### ATAC-Seq

Individual pairs of posterior forelimb fractions were dissected from 35 somite wildtype (n=2) or *Shh^-/-^* embryos (n=2). ATAC used components from the Nextera DNA Library Preparation Kit (Illumina) as described previously (Buenrostro et al. 2015) with the following variations. 5,000 cells from each sample were added into each reaction and cells were lysed on ice for 8 min. prior to centrifugation. Libraries were generated using 18 cycles of PCR amplification with NEB high fidelity 2x master mix (New England Biolabs), cleaned up with AMPure XP beads (Beckman Coulter) and sequenced on an Illumina NextSeq 500 using PEx75 to a depth of 30 million reads. Peaks were called using MACS2 with a fixed window size of 200bp and a q-value cutoff of 0.05. Differential analysis of wildtype versus *Shh^-/-^* peak signals was performed essentially as described for ChIP above using limma (Ritchie et al. 2015).

## ACKNOWLEDGMENTS

We thank Blerta Xhemalce, Samantha Brugmann, Kevin Peterson and Janani Ramachandran for comments on the manuscript. We thank Drs. Ken Zaret, Maki Iwafuchi-Doi, Jongwhan Kim and Cathy Rhee for advice on performing ATAC-seq and Jessica Podnar from the Genomic Sequencing and Analysis Facility at the University of Texas at Austin for technical advice. The Texas Advanced Computing Center (TACC) at The University of Texas at Austin provided computational resources. This work was supported by NIH R01HD073151 (to SAV and HJ), The St. Baldrick’s Foundation (to SAV) and F31DE027597 (to RKL).

## AUTHOR CONTRIBUTIONS

SV, RL, KF, ZJ and HJ conceived experiments; RL, ZJ, KF, JH and WZ performed experiments; SV, RL and KF wrote the initial draft; all authors participated in editing.

## FIGURE SUPPLEMENT LEGENDS

**Figure 1-Figure Supplement 1.**
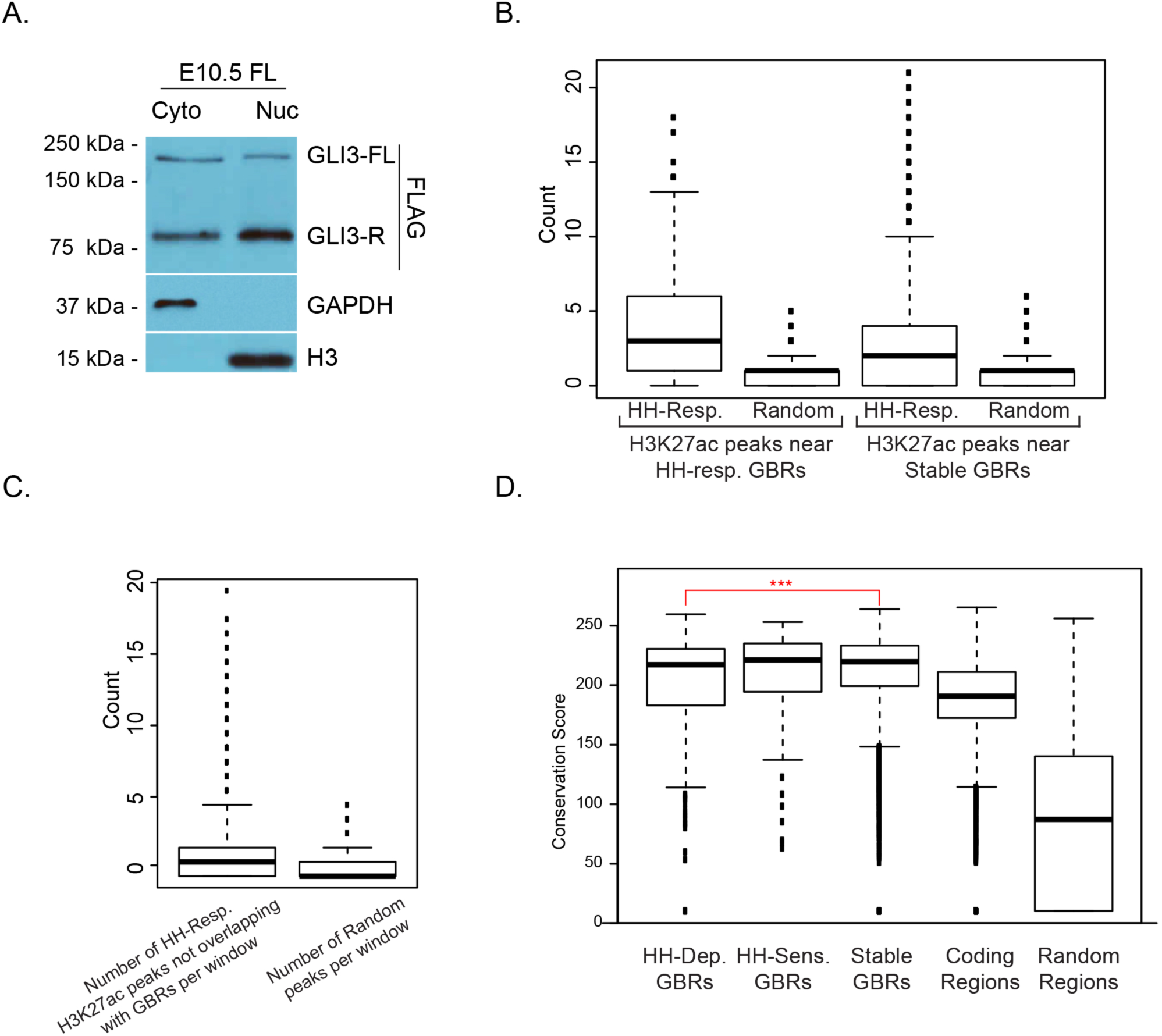
Nuclear localization of GLI3 and properties of GLI binding regions. A. Western blots from E10.5 limb buds indicating the distribution of endogenous GLI3^Flag^ in cytoplasmic and nuclear fractions. B. Hedgehog-responsive enhancers that are not bound by GLI are clustered near GLI binding regions. Box plot indicates the proximity of HH-responsive H3K27ac peaks that are not bound by GLI to either HH-Responsive GBRs or Stable GBRs compared to random peaks. For both HH-responsive and stable GBRs, the number of HH-Responsive non-GBR H3K27ac peaks is significantly larger than the number of random regions (Wilcoxon-test p-value = 0). C. HH-responsive peaks not bound by GLI3 are clustered together. The genome was split into 100,000 base-pair non-overlapping windows and the number of HH-responsive H3K27ac peaks that are not bound by GLI3 were counted as well as the number of random peaks. Only windows that overlapped with at least one HH-responsive H3K27ac peak or random peak were considered. The two counts are significantly different (Wilcoxon-test p-value = 0). The dark black line indicates the median. The lower boundary of the box indicates the first quantile, while the upper boundary of the third box is the third quantile. The circles indicate outliers. D. Box plot showing the conservation scores for different classes of GBRs. The conservation scores correspond to phastCons values linearly scaled from 0 to 255. HH-responsive GBRs have significantly lower conservation scores than stable GBRs (p-value = 0.0001134492, one sided Wilcoxon test). None of the other pairs of GBRs are significantly different from each other. ‘Coding regions’ represent conservation scores for all protein coding genes in the mouse mm10 genome while ‘Random regions’ represent conservation scores for a set of 1000 random genomic loci that do not overlap with any gene. The dark black line indicates the median. The lower boundary of the box indicates the first quantile, while the upper boundary of the third box is the third quantile. The circles indicate outliers.

**Figure 2-Figure Supplement 1.**
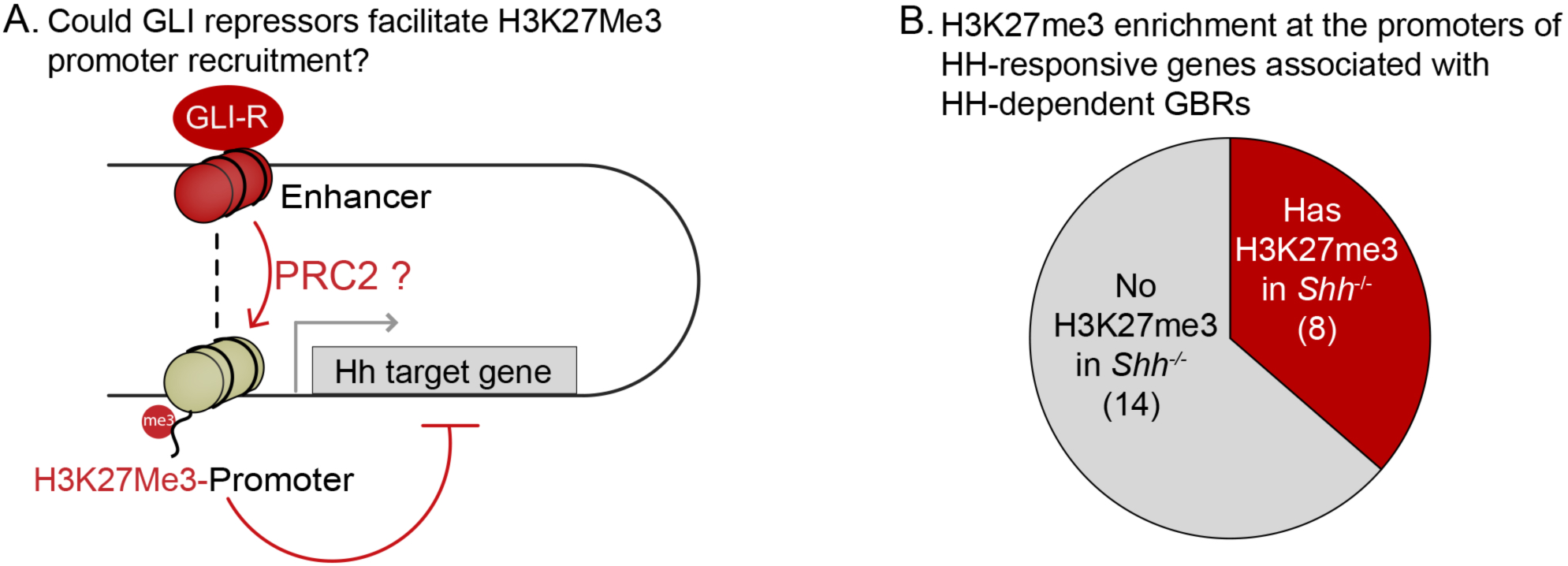
H3K27Me3 enrichment at the promoters of GLI target genes. A. Schematic illustrating a hypothetical mechanism by which GLI repressors bound to distal enhancers could facilitate the deposition of PRC2-marked H3K27Me3 at the promoters of target genes. B. H3K27Me3 enrichment within the promoters of 22 HH responsive genes that also have HH-dependent GBRs (Figure 2-Source Data 2) was determined as for the enhancers except that the reads were summed in gene promoters instead of peak regions within a window spanning from 1500 bp upstream to 500 bp downstream of the transcriptional start site. H3K27Me3 enrichment was present in the promoters of 8/22 target genes.

**Figure 4-Figure Supplement 1.**
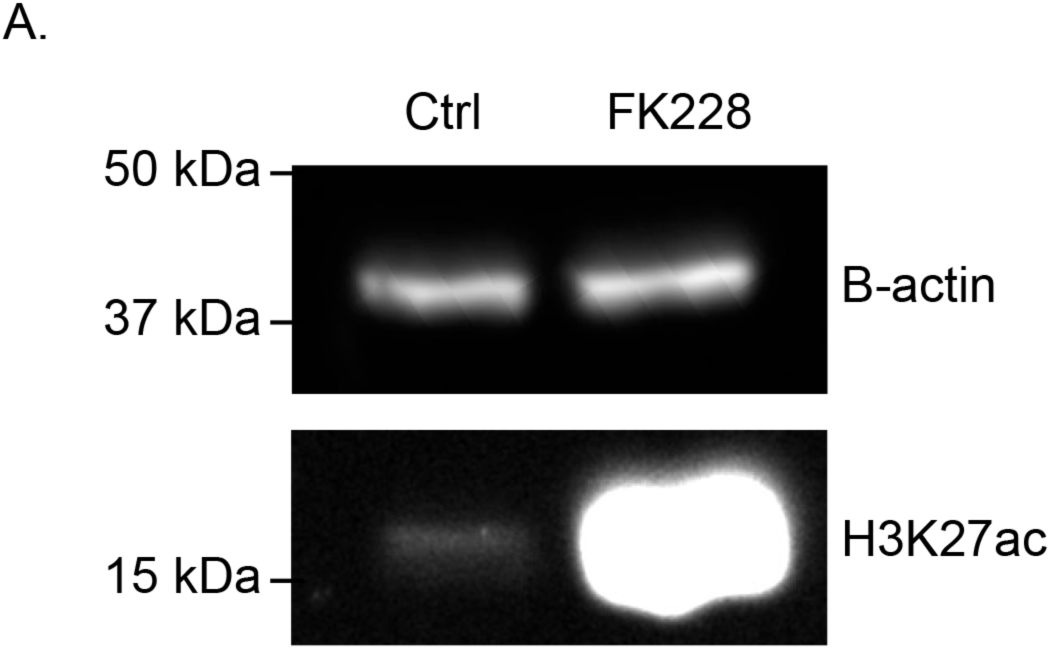
H3K27ac is increased upon HDAC inhibition. A. Western blot of cultured limb buds treated with DMSO or the HDAC1/2 inhibitor, FK228 (250nM) for 2 hours showing increased overall levels of H3K27 acetylation. Note that these are whole limb buds rather than anterior and posterior fractions shown in Figure 5.

## SOURCE DATA LEGENDS

**Figure 1-Source Data 1. Endogenous GLI3-Flag ChIP-seq analyzed data and called peaks.** GLI3 binding regions with called peaks with a false discovery rate (FDR) <0.05 from two biological replicates of E10.5 (32-35S) forelimbs. Rank ordered coordinates, peak length, log2 fold change (log2FC) and FDR are listed for each peak.

**Figure 1-Source Data 2. WT vs *Shh^-/-^* H3K27ac ChIP-seq analyzed data and called peaks.** H3K27ac called peaks with a FDR <0.05 from two biological replicates from WT and *Shh^-/-^* E10.5 forelimbs. For each peak, the assigned Peak ID, coordinates, peak type, fold change normalized to input for WT and *Shh^-/-^* samples and fold change of WT over *Shh^-/-^* are listed. Additional tabs include sorted datasets for sub-classifications. Tabs containing GBRs indicate intersections with GLI binding regions.

**Figure 1-Source Data 3. Motifs uncovered from HH-responsive enhancers**. Table showing the top 20 motifs uncovered from de novo motif analysis on HH-responsive GBRs. The enrichment is relative to matched genomic controls. Note that ‘HH_resp_2’ is the only motif with an enrichment value of greater than 2 and corresponds with a known GLI binding motif.

**Figure 2-Source Data 1. *Shh^-/-^* H3K27me3 ChIP-seq analyzed data and called peaks.** H3K27me3 called peaks with a FDR <0.05 from two replicates of *Shh^-/-^* E10.5 forelimbs. For each peak, the assigned Peak ID, coordinates, log2 fold change normalized signal to input. Additional tab includes H3K27me3 peaks that overlap with GLI3 binding regions; the GBR sub-classifications are specified.

**Figure 2-Source Data 2. Hedgehog responsive genes with H3K27me3 enrichment.** The first column indicates genes previously identified as differentially expressed between *Shh^-/-^* and WT E10.5 limb buds (Lewandowski et al. 2015). The second column indicates the fold enrichment of H3K27me3 at the promoter compared to Input with the adjusted P-value indicated in the third column. The fourth column indicates whether the gene has a HH-dependent GBR (indicated by 1 and yellow shading) within the same presumptive TAD (Dixon et al. 2012). There are 22 HH-dependent target genes out of 80 HH-responsive genes.

**Figure 2-Source Data 3. WT vs *Shh^-/-^* H3K4me2 ChIP-seq analyzed data and called peaks.** H3K4me2 called peaks with a FDR <0.05 from two replicates from WT and *Shh^-/-^* E10.5 forelimbs. For each peak, the assigned Peak ID, coordinates, peak type, fold change normalized to input for WT and *Shh^-/-^* samples and fold change of WT over *Shh^-/-^* are listed. Additional tabs include sorted files for each peak type. Under the ‘GLI3 binding’ column, ‘TRUE’ implies overlap with a GBR, while ‘FALSE’ indicates no overlap.

**Figure 3-Source Data 1. WT vs *Shh^-/-^* ATAC-Seq analyzed data and called peaks.** Coordinates for all ATAC peaks in the WT group that overlap with GBRs are listed. “Shh_ATAC_peak” identifies the corresponding id# for that peak in the *Shh^-/-^* data, and if a peak is not present in the *Shh^-/-^* samples, it is marked as NA. A column for each GBR type identifies which GBR type a given ATAC peak overlaps with. The number indicates the peak ID. If a peak region does not overlap with the type of peak in that list, it will be marked as NA. The normalized log2 transformed signals are showed for each sample in addition to the “average” signal across all samples. The “t” statistic calculates the difference in signals between WT and *Shh^-/-^* by taking into consideration fold-change and variance among samples. A positive t statistic values indicate a peak is more accessible in WT than *Shh^-/-^* and a negative t statistic indicates higher accessibility in *Shh^-/-^*. The “p.value” is obtained from a moderated t-test using limma. The “p.value.adj” is the adjusted p-value (FDR) using the Benjamini-Hochberg procedure.

**Figure 4-Source Data 1. WT vs *Shh^-/-^;Gli3^-/-^* H3K27ac MicroChIP-seq analyzed data and called peaks.** H3K27ac called peaks with a FDR <0.05 from two replicates of WT, *Shh^-/-^* and *Shh^-/-^;Gli3^-/-^* E10.5 (33-34S) forelimbs. Separate tabs for each genotype include peak coordinates and log2 fold change over input. Additional tabs include a peak summary and differential analysis of WT vs. *Shh^-/-^;Gli3^-/-^*. Differential analysis tab lists peak coordinates, peak type, fold change normalized to input for WT and *Shh^-/-^;Gli3^-/-^* samples and fold change of WT over *Shh^-/-^;Gli3^-/-^*.

**Figure 5-Source Data 1. Stable and HH-responsive GLI binding regions with limb enhancer activity in the VISTA dataset.** Columns indicate VISTA enhancer IDs, coordinates, number of tissues with limb enhancer activity as annotated by the VISTA database (Visel et al. 2007) and its corresponding GBR category.

**Figure 5-Source Data 2. NIH3T3 H3K27ac ChIP-seq analyzed data and called peaks.** H3K27ac called peaks with a FDR <0.05 from two replicates of purmorphamine (“pm”) treated or DMSO control NIH3T3 cells. For each peak, the assigned Peak ID, coordinates, peak type, fold change normalized to input for purmorphamine treated and control samples, and fold change of purmorphamine treated over control are listed. Additional tabs include sorted files for each peak type.

